# Muscarinic receptor activation preferentially inhibits rebound in vulnerable dopaminergic neurons

**DOI:** 10.1101/2024.07.30.605819

**Authors:** Megan L. Beaver, Rebekah C. Evans

## Abstract

Dopaminergic subpopulations of the substantia nigra *pars compacta* (SNc) differentially degenerate in Parkinson’s disease and are characterized by unique electrophysiological properties. The vulnerable population expresses a T-type calcium channel-mediated afterdepolarization (ADP) and shows rebound activity upon release from inhibition, whereas the resilient population does not have an ADP and is slower to fire after hyperpolarization. This rebound activity can trigger dopamine release in the striatum, an important component of basal ganglia function. Using whole-cell patch clamp electrophysiology on *ex vivo* slices from adult mice of both sexes, we find that muscarinic activation with the non-selective muscarinic agonist Oxotremorine inhibits rebound activity more strongly in vulnerable vs resilient SNc neurons. Here, we show that this effect depends on the direct activation of muscarinic receptors on the SNc dopaminergic neurons. Through a series of pharmacological and transgenic knock-out experiments, we tested whether the muscarinic inhibition of rebound was mediated through the canonical rebound-related ion channels: T-type calcium channels, hyperpolarization-activated cation channels (HCN), and A-type potassium channels. We find that muscarinic receptor activation inhibits HCN-mediated current (I_h_) in vulnerable SNc neurons, but that I_h_ activity is not necessary for the muscarinic inhibition of rebound activity. Similarly, we find that Oxotremorine inhibits rebound activity independently of T-type calcium channels and A-type potassium channels. Together these findings reveal new principles governing acetylcholine and dopamine interactions, showing that muscarinic receptors directly affect SNc rebound activity in the midbrain at the somatodendritic level and differentially modify information processing in distinct SNc subpopulations.

**Significance Statement:** Dopaminergic neurons in the substantia nigra *pars compacta* (SNc) can be divided into functional subpopulations with distinct basal ganglia connectivity and different degeneration patterns in Parkinson’s disease. We show that the vulnerable and resilient subpopulations of SNc dopaminergic neurons are differentially modulated by muscarinic receptor activation. Specifically, muscarinic receptor activation inhibits rebound activity more strongly in the vulnerable SNc neurons than in the resilient. We find that this inhibition occurs through a non-canonical rebound-related pathway and is not mediated through the channels best known for modulating rebound in midbrain dopaminergic neurons. These findings are important because they reveal novel acetylcholine-dopamine interactions that occur in the midbrain and affect information processing in distinct basal ganglia circuits.

## Introduction

Dopaminergic neurons of the midbrain play a significant role in behaviors including aversion, reward learning, and voluntary movement. Degeneration of the dopamine neurons of the substantia nigra *pars compacta* (SNc) is responsible for many of the symptoms associated with Parkinson’s disease (PD). However, SNc neurons do not degenerate uniformly. Two populations of cells, mapped along the dorsal-ventral axis, can also be defined by their vulnerability or resilience to degeneration in PD (Yamada et al., 1990; Fearnley and Lees, 1991; Gibb and Lees, 1991; Damier et al., 1999). These ventral and dorsal tier populations are involved in different basal ganglia circuits (Evans et al., 2020) and process information in distinct ways (Evans et al., 2017). The ventral tier, which is more prone to degeneration, contains dopaminergic neurons that express aldehyde dehydrogenase 1a1 but not calbindin (Poulin et al., 2014, 2020; Wu et al., 2019; Carmichael et al., 2021). These vulnerable neurons also have a distinct electrophysiological signature in that they display an afterdepolarization (ADP) when activated from hyperpolarized potentials (Evans et al., 2017). This ADP is mediated by a high number of T-type calcium channels, which in combination with a large number of hyperpolarization-activated cation (HCN) channels enhance rebound firing (Neuhoff et al., 2002; Evans et al., 2017). Previous studies have identified rebound activity as a mode of specialized dopaminergic information processing that is unique to the ventral tier of the SNc and has been observed both *in vivo* and in *ex vivo* slices (Fiorillo et al., 2013b; Evans et al., 2020).

Multiple ion channels are responsible for mediating the rebound response, including T-type calcium channels, HCN channels, and A-type potassium channels. T-type calcium and HCN channels, which are activated at hyperpolarized potentials, work to rapidly depolarize the cell after inhibition is released (Mercuri et al., 1995; Neuhoff et al., 2002; Amendola et al., 2012; Evans et al., 2017). This rapid depolarization is countered by A-type potassium channels whose outward current prolongs the rebound delay as they inactivate (Tarfa et al., 2017). These three channel types can be actively suppressed or enhanced by neuromodulators (Hildebrand et al., 2007; Gambardella et al., 2012; Gantz and Bean, 2017), suggesting that rebound activity in dopamine neurons is a dynamically modulated characteristic.

One neuromodulator, acetylcholine, is of particular interest as it is known to have important interactions with the dopaminergic system. Extensive previous literature details the influence of striatal cholinergic interneurons over dopamine release in the striatum (Zhou et al., 2001; Zhang and Sulzer, 2004; Pakhotin and Bracci, 2007; Threlfell et al., 2010; Nelson et al., 2014; Shin et al., 2015; Kramer et al., 2022; Razidlo et al., 2022; Krok et al., 2023). However, less is known about the influence of acetylcholine, especially muscarinic receptor activation, on dopaminergic cell bodies and dendrites in the midbrain. Most dopaminergic neurons in the SNc, regardless of subtype, express M5 muscarinic receptors (Weiner et al., 1990). These are G_q_-protein-coupled receptors that have been shown to increase intracellular Ca_2+_ in SNc dopamine neurons (Foster et al., 2014) and alter action potential characteristics (Scroggs et al., 2001). *In vivo*, muscarinic receptors in the midbrain mediate long-lasting dopamine release in the striatum (Forster and Blaha, 2003; Steidl et al., 2011) and M5-specific modulation alters effort-choice and depression-related behaviors (Nunes et al., 2020, 2023). However, the influence that these M5 receptors have on the intrinsic properties and rebound activity of the different SNc subpopulations has not been comprehensively investigated.

Here we combine whole-cell patch clamp electrophysiology and pharmacology to evaluate the effects of muscarinic receptor activation on SNc subpopulations. We find that muscarinic activation strongly reduces rebound activity in the vulnerable SNc neural subtype, but only weakly reduces it in the more resilient SNc neural subtype. By selectively blocking the channels known to mediate SNc rebound activity, we show that muscarinic activation of SNc neurons inhibits rebound activity through a non-canonical mechanism.

## Materials and Methods

### Animal use

All animal handling and procedures were approved by the Animal Care and Use Committee for Georgetown University. *Dopamine transporter (DAT)-cre/Ai9* mice [B6.SJL-*Slc6a3*_*tm1*.*1(cre)Bkmn*_/J, JAX #006660, (Bäckman et al., 2006) /B6.Cg-*Gt(ROSA)26Sor*_*tm9(CAG-tdTomato)Hze*_/J, JAX #007909, (Madisen et al., 2010)] of either sex were used at age postnatal day >60 (average age: 120 ± 9 days). CaV3.3 KO mice [*Cacna1i*^*-/-*^ on C57BL/6J background; Courtesy of Broad Institute of MIT and Harvard, (Ghoshal et al., 2020)], were used where specified (average age: 41 ± 3 days).

### Slice preparation

Mice were anesthetized with inhaled isoflurane and transcardially perfused with an ice-cold, oxygenated, glycerol-based modified artificial cerebrospinal fluid (aCSF) solution containing the following (in millimolar (mM)): 198 glycerol, 2.5 KCl, 1.2 NaH_2_PO_4_, 25 NaHCO_3_, 20 HEPES, 10 glucose, 10 MgCl_2_, and 0.5 CaCl_2_. Mice were then decapitated and brains extracted. Coronal midbrain slices (200 µm) containing the substantia nigra region were prepared using a vibratome (Leica VT 1200S) and incubated for 30 minutes in heated (34°C) oxygenated holding aCSF containing (in mM): 92 NaCl, 30 NaHCO_3_, 1.2 NaH_2_PO_4_, 2.5 KCl, 35 glucose, 20 HEPES, 2 MgCl_2_, 2 CaCl_2_, 5 Na-ascorbate, 3 Na-pyruvate, and 2 thiourea as in Evans et al., 2017. Slices, in their holding chamber, were incubated at room temperature for at least 30 mins.

### Electrophysiological recordings

Slices were hemisectioned and continuously superfused at ∼2 mL/min with warm (34°C), oxygenated extracellular recording solution containing the following (in mM): 125 NaCl, 25 NaHCO_3_, 3.5 KCl, 1.25 NaH_2_PO_4_, 10 glucose, 1 MgCl_2_, and 2 CaCl_2_. Neurons were visualized with a 40x objective using a Prior OpenStand Olympus microscope equipped with a scientific CMOS camera (Hamamatsu ORCA-spark).

Whole-cell recordings were made using borosilicate pipettes (2-5 MΩ) pulled with a flaming/brown micropipette puller (Sutter Instruments) and filled with internal recording solution containing (in mM): 121.5 KMeSO_3_, 9 NaCl, 9 HEPES, 1.8 MgCl_2_, 14 phosphocreatine, 4 Mg-ATP, 0.3 Na-GTP, 0.1 CaCl_2_, and 0.5 EGTA adjusted to a pH value of ∼7.35 with KOH. All salts were purchased from Sigma-Aldrich.

Signals were digitized with an Axon Digidata 1550B interface, amplified by a Multiclamp 700B amplifier, and acquired using Clampex11.2 software (Molecular Devices). Data were sampled in current clamp at 10 kHz and in voltage clamp at 100kHz with filtering at 5 kHz. Data were analyzed using custom procedures in Igor Pro (WaveMetrics).

All recordings were performed in dopaminergic neurons which were targeted by their anatomic location and presence of TdTomato, where applicable, and identified based on various electrophysiological characteristics, such as the firing frequency (<5 Hz) and presence of HCN-mediated sag. Ventral-tier (vulnerable) SNc neurons were identified by the presence of the distinctive ADP (Evans et al., 2017). Each slice was used for only one drug wash-on series (one cell).

### Drugs

Patch-clamp recordings were performed in the presence of synaptic blockers [10 µM Gabazine (Tocris Bioscience), 1 µM CGP-35348 (Tocris Bioscience), 5 µM NBQX (Tocris Bioscience), and 50 µM D-AP5 (Hello Bio)], unless otherwise specified. As indicated, we used 3 µM Oxotremorine (Sigma-Aldrich), 10 µM Atropine (Sigma-Aldrich), 1 µM TTA-P2 (Alomone Labs), 10 µM ZD7288 (Hello Bio), and/or 100 nM AmmTx3 (Alomone Labs). All drugs were prepared as aliquots in water or DMSO.

### Data analysis

Data were analyzed using Igor Pro (WaveMetrics) and GraphPad Prism. Statistical significance in two group comparisons was determined using Wilcoxon rank-sum tests (unpaired) or Wilcoxon signed-rank tests (paired). Statistical significance in three group comparisons was determined using Kruskal-Wallis tests followed by Dunn’s multiple comparisons tests, where applicable. Descriptive statistics are reported as mean ± standard error of the mean (SEM) and error shading on graphs indicates ± SEM. Box plots show median, 25^th^ and 75^th^ percentiles (boxes), and 9^th^ and 91^st^ percentiles (whiskers). For each treatment group, n indicates number of cells, with no more than 1 cell per slice or 3 cells per treatment condition from a single mouse.

We evaluated rebound using several different measures in order to provide a comprehensive understanding in the changes in rebound activity elicited by muscarinic receptor activation. We recorded rebound and the ADP in a current-clamp protocol that hyperpolarizes the cell to approximately -80 mV, stimulates a single action potential from hyperpolarization, and then releases the hyperpolarization (Figure 1A). We determined the rebound slope to be the most reliable measure of rebound, measured as the slope of depolarization to the first action potential when released from hyperpolarization. Next, we measured the rebound delay – the time it takes the cell to fire an action potential once released from hyperpolarization. If a cell did not fire an action potential within 1 second of repolarization, the rebound delay was recorded as 1 second. Not all cells consistently fire action potentials during the rebound period. However, these cells do show a characteristic and measurable depolarizing slope, even if they do not reach threshold to fire an action potential. Finally, when possible, we measured rebound frequency as the frequency of the first two spikes upon release from hyperpolarization. If the first two spikes do not occur within the 1 second rebound period or there is only one spike, the rebound frequency was recorded as zero.

**Figure 1.**
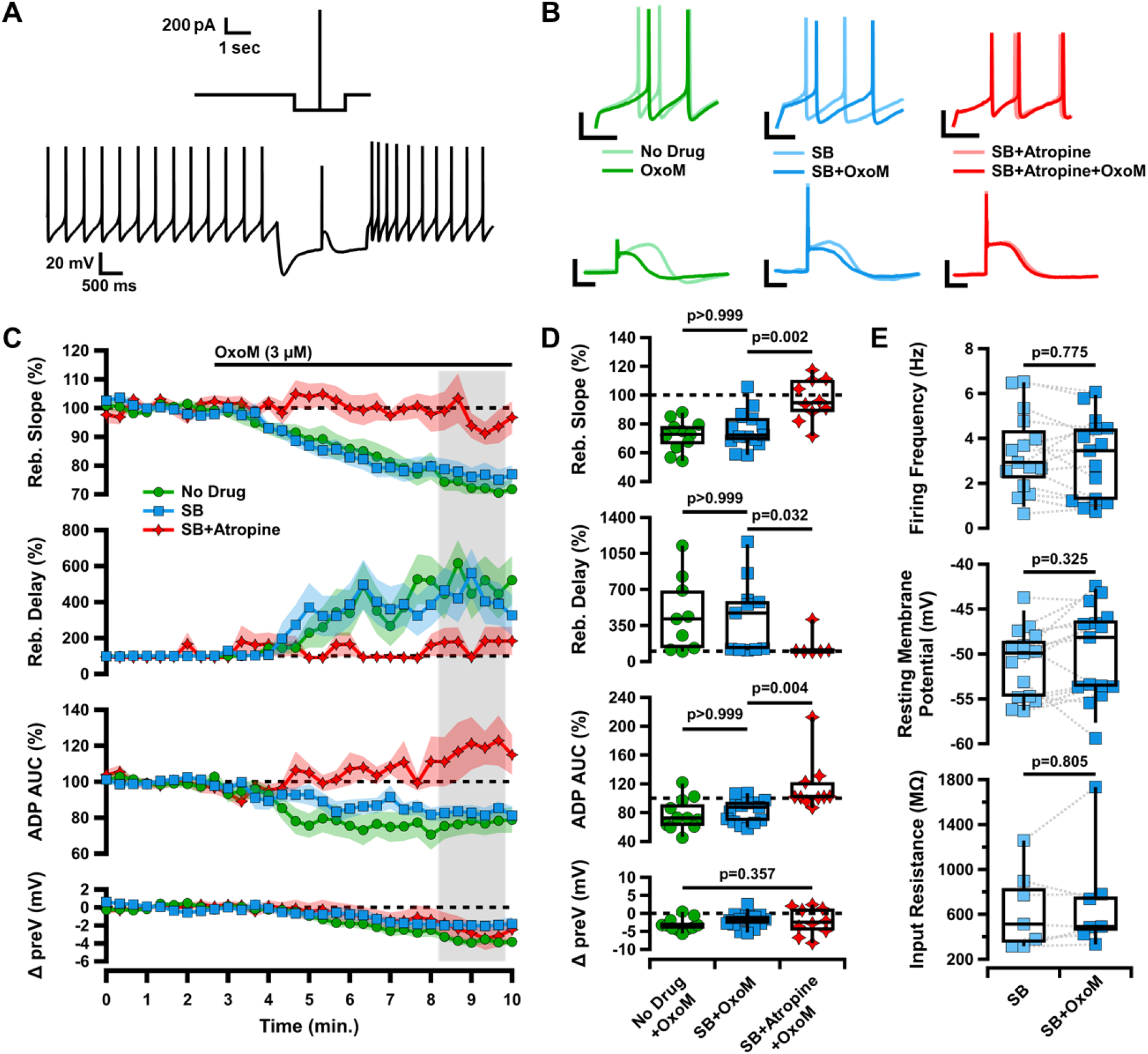
Oxotremorine (OxoM) inhibits rebound of SNc neurons through post-synaptic muscarinic receptors. **A**, Diagram of current-clamp protocol used to elicit rebound and the ADP (top). Sample trace of entire current clamp protocol (bottom). **B**, Sample traces of rebound (top) and the ADP (bottom) before (light) and after (dark) application of OxoM with no drug (green), synaptic blockers (SB) (blue), or SB+Atropine (red) in the bath solution. Scale bars: 20 mV, 100 ms. **C**, Normalized rebound slope (top), rebound delay (second), ADP area under the curve (AUC) (third), and hyperpolarized baseline (bottom) as a function of time. Data presented as average ±SEM. **D**, Box plots representing individual cell averages in shaded regions of C. There were significant differences between groups in rebound slope (p=0.002, Kruskal-Wallis), rebound delay (p=0.020, Kruskal-Wallis), and ADP (p=0.004, Kruskal-Wallis), but not hyperpolarized baseline (p=0.357, Kruskal-Wallis). Remaining p values are from Dunn’s test. **E**, Box plots showing intrinsic characteristics of cells in SB before (light blue) and after (dark blue) application of OxoM.

### Results

### Oxotremorine inhibits rebound of SNc neurons through post-synaptic muscarinic receptors

To investigate the effects of muscarinic acetylcholine receptor (mAChR) activation on SNc neurons, we performed whole-cell patch clamp electrophysiology on coronal slices from DAT-cre/Ai9 mice. In these slices, dopaminergic SNc neurons were identified by their red fluorescence and divided into subpopulations based on the presence or absence of an electrophysiologically-recorded calcium-mediated afterdepolarization (ADP) when stimulated from a hyperpolarized potential, as in Evans et al., 2017 (Figure 1A). During current-clamp recordings of each dopaminergic SNc neuron, we washed on 3 µM Oxotremorine (OxoM), a non-selective muscarinic agonist, to activate mAChRs. We found that OxoM reliably decreased the rebound activity of ADP-expressing SNc neurons (Figure 1C). We evaluated the effect of muscarinic activation on SNc neuron characteristics in three ways: 1. rebound slope (72.07±3.25% of baseline due to OxoM), 2. rebound delay (454.23±119.97% of baseline due to OxoM), and 3. area under the curve (AUC) of the ADP (77.45±6.75% of baseline due to OxoM). The ADP is elicited by stimulating an action potential from a hyperpolarized potential (reaching approximately - 80 mV). Rebound slope and rebound delay are measured when releasing the cell from a hyperpolarized potential (Figure 1A, see methods for details).

To test whether OxoM inhibited dopaminergic rebound activity by altering presynaptic glutamatergic or GABAergic inputs to the recorded cell, we applied OxoM in the presence of synaptic blockers (SB; 10 µM gabazine, 1 µM CGP-35348, 5 µM NBQX, and 50 µM AP5) (Figure 1C-D). In both the presence and absence of synaptic blockers, OxoM consistently reduced rebound slope [no drug (ND) n=11, 72.07±3.25% ; SB n=15, 76.34±3.30%; Dunn’s p>0.999], increased rebound delay (ND n=9, 454.23±119.97%; SB n=11, 437.65±107.74%; Dunn’s p>0.999), and reduced ADP size (ND n=9, 77.45±6.75%; SB n=11, 83.19±4.24%; Dunn’s p>0.999). These findings show that the effect of OxoM on dopaminergic rebound properties is a direct effect on the post-synaptic SNc neuron, rather than an effect on pre-synaptic neurotransmitter release.

To determine whether the OxoM-mediated rebound inhibition required post-synaptic muscarinic receptor activation, we applied OxoM in the presence of the mAChR antagonist Atropine (10 µM) (Figure 1C-D). In the presence of Atropine (Atp.), OxoM did not reduce rebound slope (SB n=15, 76.34±3.30%; SB+Atp. n=10, 96.59±4.55%; Dunn’s p=0.002), rebound delay (SB n=11, 437.65±107.74%; SB+Atp. n=6, 152.09±51.35%; Dunn’s p=0.032), or ADP size (SB n=15, 83.19±4.24%; SB+Atp. n=10, 115.61±11.50%, Dunn’s p=0.004). This finding shows that OxoM reduces rebound properties through activation of muscarinic acetylcholine receptors.

There was no significant difference in the change in hyperpolarized membrane potential (preV) between OxoM alone or OxoM with synaptic blockers or synaptic blockers and Atropine (p=0.357, Kruskal-Wallis). Further, there was no significant change in spontaneous activity of these cells in synaptic blockers with the application of OxoM (Figure 1E), as measured by firing frequency (n=15; SB 3.37±0.46 Hz, SB+OxoM 3.14±0.47 Hz; Wilcoxon signed-rank p=0.775), resting membrane potential (n=15; SB - 51.01±1.00 mV, SB+OxoM -49.84±1.35 mV; Wilcoxon signed-rank p=0.325), and input resistance (n=7; SB 635.83±134.50 MΩ, SB+OxoM 712.34±181.16 MΩ; Wilcoxon signed-rank p=0.805). Together, these results indicate that OxoM reduces dopaminergic rebound activity through actions of post-synaptic muscarinic receptors.

### Inhibition of rebound by OxoM is strongest in ventral-tier SNc neurons

The population of neurons in the ventral tier of the SNc has a higher expression of T-type calcium channels (TTCCs) and HCN channels (Mercuri et al., 1995; Neuhoff et al., 2002; Evans et al., 2017). As a result, these neurons demonstrate the TTCC-mediated ADP and fire faster during rebound than their dorsal tier non-ADP counterparts (Figure 2A). We wanted to determine if the inhibition of rebound by OxoM was unique to ADP cells, or if rebound firing in non-ADP cells was also inhibited by mAChR activation. We found that OxoM application had a significantly stronger effect on the rebound activity of ADP cells compared to non-ADP cells. OxoM application to non-ADP cells resulted in a smaller decrease in rebound slope (ADP with SB n=15, 76.34±3.30%; non-ADP with SB n=16, 88.16±3.45%; Wilcoxon rank-sum p=0.011) and rebound frequency (ADP with SB n=10, 19.61±11.19%; non-ADP with SB n=16, 63.42±8.61%; Wilcoxon rank-sum p=0.024) as compared to ADP cells (Figure 2C-D). Further, they showed a smaller increase in rebound delay (ADP with SB n=11, 437.65±107.74; non-ADP with SB n=16, 173.80±41.79%; Wilcoxon rank-sum p=0.013) as compared to ADP cells. No difference was observed in the change in hyperpolarized baseline between groups (ADP with SB n=15, -1.94±0.54 mV; non-ADP with SB n=16, -2.76±0.47 mV; Wilcoxon rank-sum p=0.216). As observed in ADP cells, OxoM did not affect other intrinsic properties of the non-ADP cells (*data not shown*). Therefore, we conclude that mAChR activation strongly inhibits rebound in ADP cells, but only weakly affects rebound in non-ADP cells.

**Figure 2.**
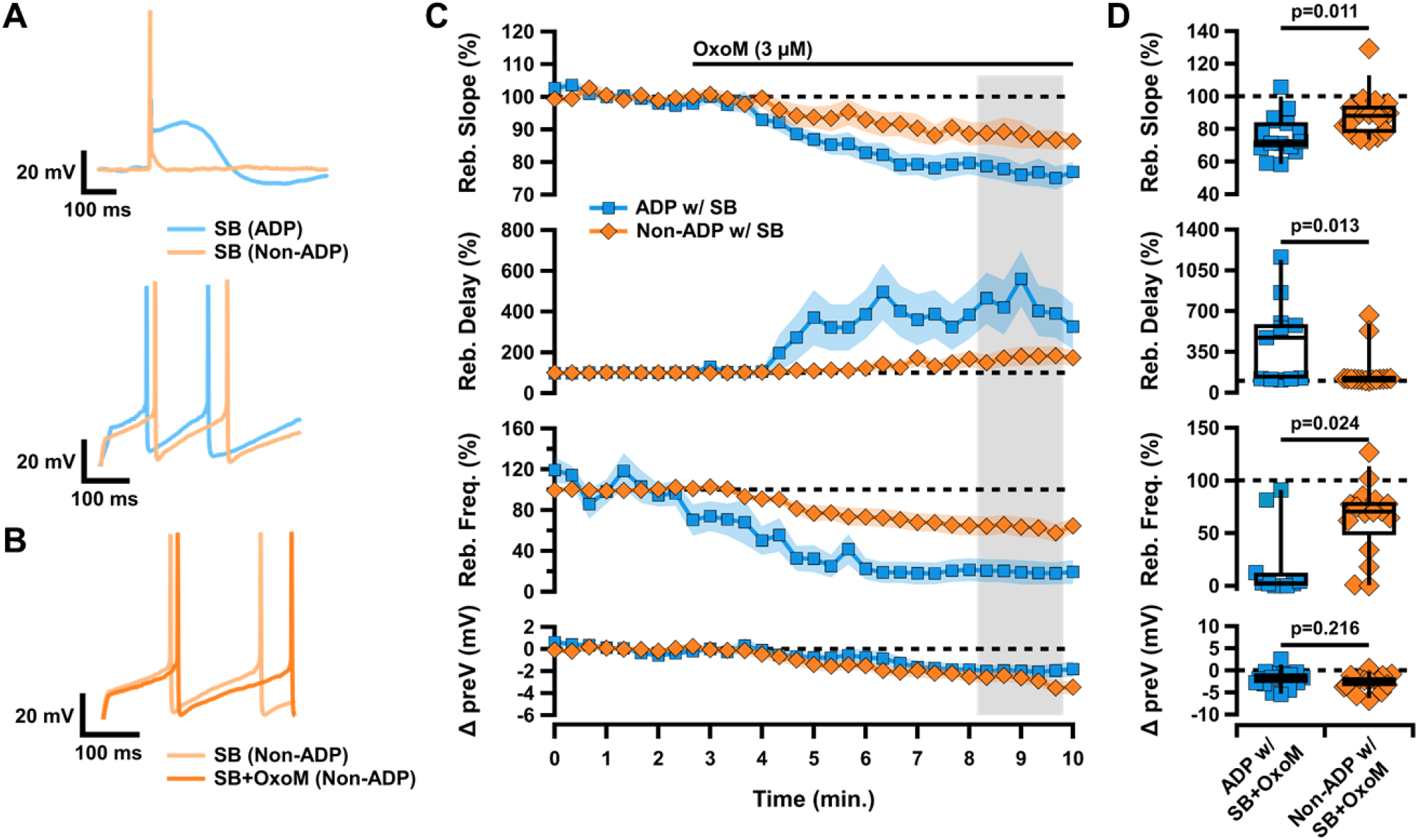
Muscarinic activation differentially inhibits rebound in SNc subpopulations. **A**, Sample traces of ADP (blue) vs non-ADP (orange) cells showing an action potential elicited from a hyperpolarized baseline (top) and rebound after release from hyperpolarization (bottom). **B**, Sample trace of rebound in a non-ADP cell in SB before (light orange) and after (dark orange) application of OxoM. **C**, Normalized rebound slope (top) rebound delay (second), rebound frequency (middle), and hyperpolarized baseline (bottom) as a function of time. Data presented as average ±SEM. **D**, Box plots representing individual cell averages in shaded regions of C.

### OxoM inhibits rebound independent of T-type calcium channels

The ventrally-located cells of the SNc contain large amounts of TTCCs which mediate rebound firing and the ADP (Evans et al., 2017). Dopaminergic SNc neurons selectively express the M5 muscarinic receptor which is G_q_-coupled (Weiner et al., 1990; Offermanns et al., 1994; Caulfield and Birdsall, 1998). Interestingly, G_q_-coupled muscarinic receptors have been shown to inhibit TTCCs in cultured cells (Hildebrand et al., 2007). Because mAChR activation inhibits rebound in the ADP cells more strongly than in the non-ADP cells, we hypothesized that OxoM inhibits rebound activity by inhibiting TTCCs. To test this, we applied OxoM in the presence of TTA-P2 (1 µM), a pan-TTCC blocker. On its own, TTA-P2 completely eliminated the ADP and reduced rebound activity (*data not shown*), as demonstrated previously (Evans et al., 2017). Surprisingly, however, the presence of TTA-P2 did not occlude the inhibitory effect of OxoM on rebound activity (Figure 3A-B). In the presence of TTA-P2, there was no significant difference in the effect of OxoM on rebound slope (SB n=15, 76.34±3.30%; SB+TTA-P2 n=10, 78.12±3.67%; Wilcoxon rank-sum p=0.765), rebound delay (SB n=11, SB+TTA-P2 n=8; Wilcoxon rank-sum p=0.492), rebound frequency (SB n=10, 19.61±11.19%; SB+TTA-P2 n=7, 48.18±14.43%; Wilcoxon rank-sum p=0.113), or hyperpolarized baseline (SB n=15, - 1.94±0.54 mV; SB+TTA-P2 n=10, -1.68±0.46; Wilcoxon rank-sum p=0.807). Though not statistically significant, there was a slight reduction in the effect of OxoM on rebound delay and frequency in the presence of TTA-P2 (Figure 3A-B).

**Figure 3.**
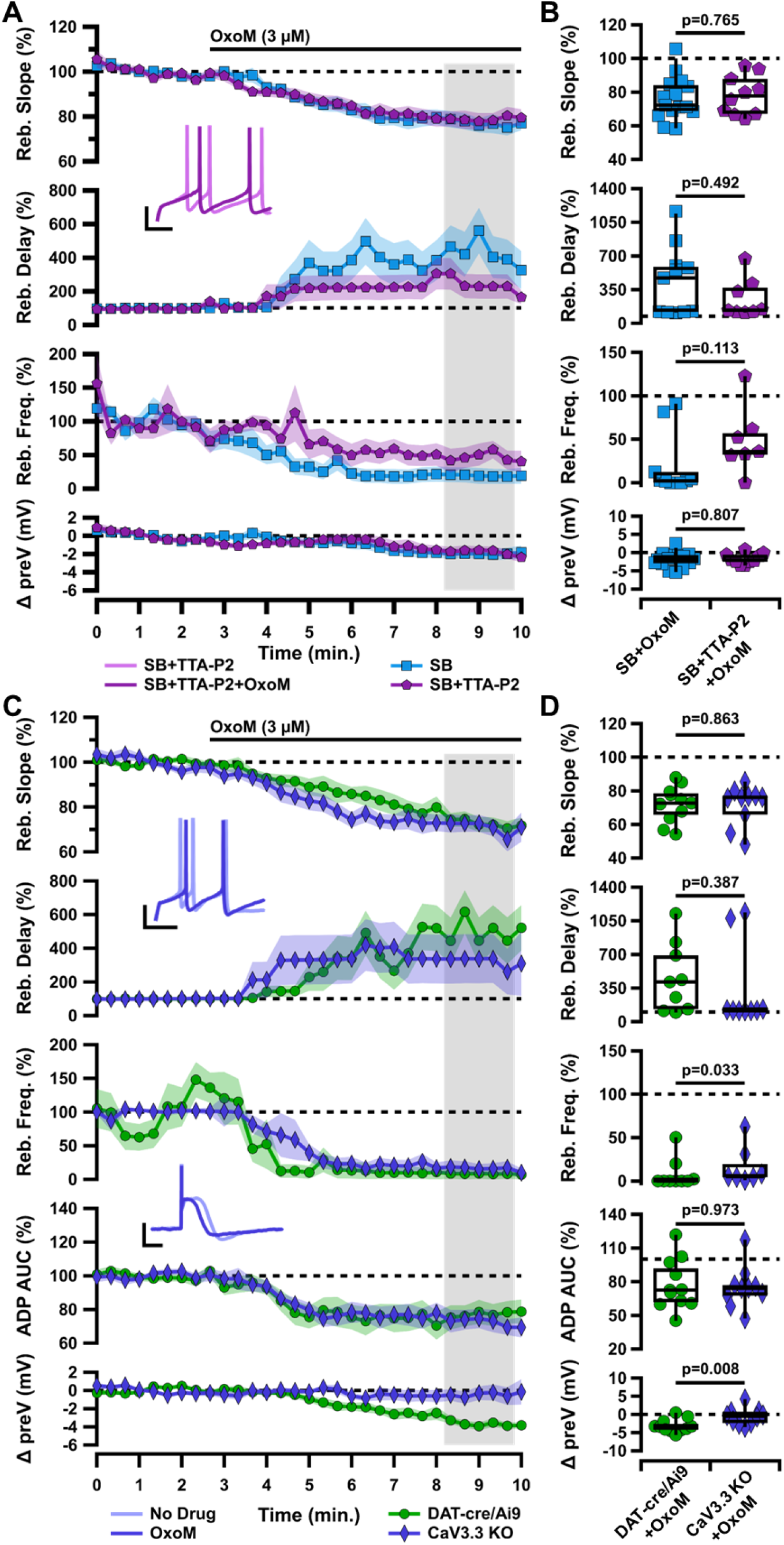
Muscarinic inhibition of rebound and the ADP of ventral tier SNc neurons is not mediated by T-type calcium channels. **A**, Normalized rebound slope (top), rebound delay (second), rebound frequency (third), and hyperpolarized baseline (bottom) as a function of time. Data presented as average ±SEM. Inset: Sample traces of rebound in SB+TTA-P2 before (light purple) and after (dark purple) application of OxoM (bottom). Scale bars: 20 mV, 100 ms. **B**, Box plots representing individual cell averages in shaded regions of A. **C**, Normalized rebound slope (top), rebound delay (second), rebound frequency (third), ADP AUC (fourth), and hyperpolarized baseline (bottom) as a function of time. Data presented as average ±SEM. Inset: Sample traces of rebound (top) and the ADP (bottom) before (light indigo) and after (dark indigo) application of OxoM in CaV3.3 KO mice. Scale bars: 20 mV, 100 ms. **D**, Box plots representing individual cell averages in shaded regions of C.

Because TTA-P2 completely abolished the ADP, we were not able to use ADP size as a measure in these experiments. As ADP size would be the measure most sensitive to an OxoM effect on TTCCs, we decided to investigate whether OxoM may be selectively inhibiting one subtype of TTCC. Of the three members of the TTCC family, CaV3.3 channels display slower activation and inactivation kinetics than the CaV3.1 and CaV3.2 subtypes (McRory et al., 2001; Chemin et al., 2002). Though the presence of CaV3.3 in SNc neurons is controversial (Dryanovski et al., 2013; Dufour et al., 2014; Poetschke et al., 2015; Guzman et al., 2018; Benkert et al., 2019), there is clear evidence that mAChRs (G_q_-coupled) can inhibit CaV3.3 in cultured cells (Hildebrand et al., 2007). We hypothesized that selective inhibition of CaV3.3 by OxoM could be responsible for the observed decrease in rebound and ADP size. Using a CaV3.3 knockout mouse (*Cacna1i*^*-/-*^ (Ghoshal et al., 2020), we performed the same electrophysiology experiments with application of OxoM. We observed no difference in ADP size at baseline between knockout and wild-type conditions (*data not shown*), allowing us to test the effect of OxoM on ADP size in these experiments. We found that OxoM reduced rebound activity similarly in ADP cells in both CaV3.3 KO and DAT-cre/Ai9 mice (Figure 3C-D). These experiments were performed without synaptic blockers in the bath solution. There was no difference between KO and DAT-cre/Ai9 mice in the OxoM reduction of rebound slope (DAT n=11, 72.07±3.25%; KO n=10, 71.60±3.73%; Wilcoxon rank-sum p=0.863), enhancement of rebound delay (DAT n=9, 454.23±119.97%; KO n=9, 338.41±145.31%; Wilcoxon rank-sum p=0.387), or reduction of ADP size (DAT n=11, 77.45±6.75%; KO n=10, 74.83±5.87%; Wilcoxon rank-sum p=0.973). Surprisingly, OxoM did not lower the hyperpolarized baseline in the KO animals as it did in DAT-cre/Ai9 (DAT n=11, -3.13±0.55 mV; KO n=10, -0.55±0.71 mV; Wilcoxon rank-sum p=0.008). While the mechanism underlying this difference is not clear, these results show that the effect of OxoM on rebound measures is not related to or dependent on OxoM’s slight augmentation of the hyperpolarized baseline membrane potential. Therefore, from this set of experiments, we concluded that OxoM’s effect on rebound is not mediated by TTCCs.

### OxoM inhibition of HCN channels is not the mechanism of reduced rebound

The hyperpolarization-activated cation current (I_h_) is mediated by HCN channels and plays a role in rebound firing, as it is activated by hyperpolarization and slow to turn off following return to resting membrane potential when released from inhibition (Mercuri et al., 1995; Neuhoff et al., 2002). In voltage-clamp recordings before and after application of OxoM, we find that I_h_ is inhibited by mAChR activation (Figure 4A-B). Cells were held at -60 mV and I_h_ currents were elicited with 1 second voltage steps (ranging -50 mV to - 120 mV in 5 mV increments) followed by a 500 ms voltage step to -120 mV to measure the tail currents. Normalized tail current amplitude was plotted as the function of the test potentials and fitted with the Boltzmann equation (Figure 4B). There was a significant decrease in the voltage for half-maximal activation (V50) of I_h_ in control vs OxoM conditions (Figure 4C; n=7, SB -97.25±1.30 mV, SB+OxoM -102.55±1.58; Wilcoxon signed-rank p=0.016).

**Figure 4.**
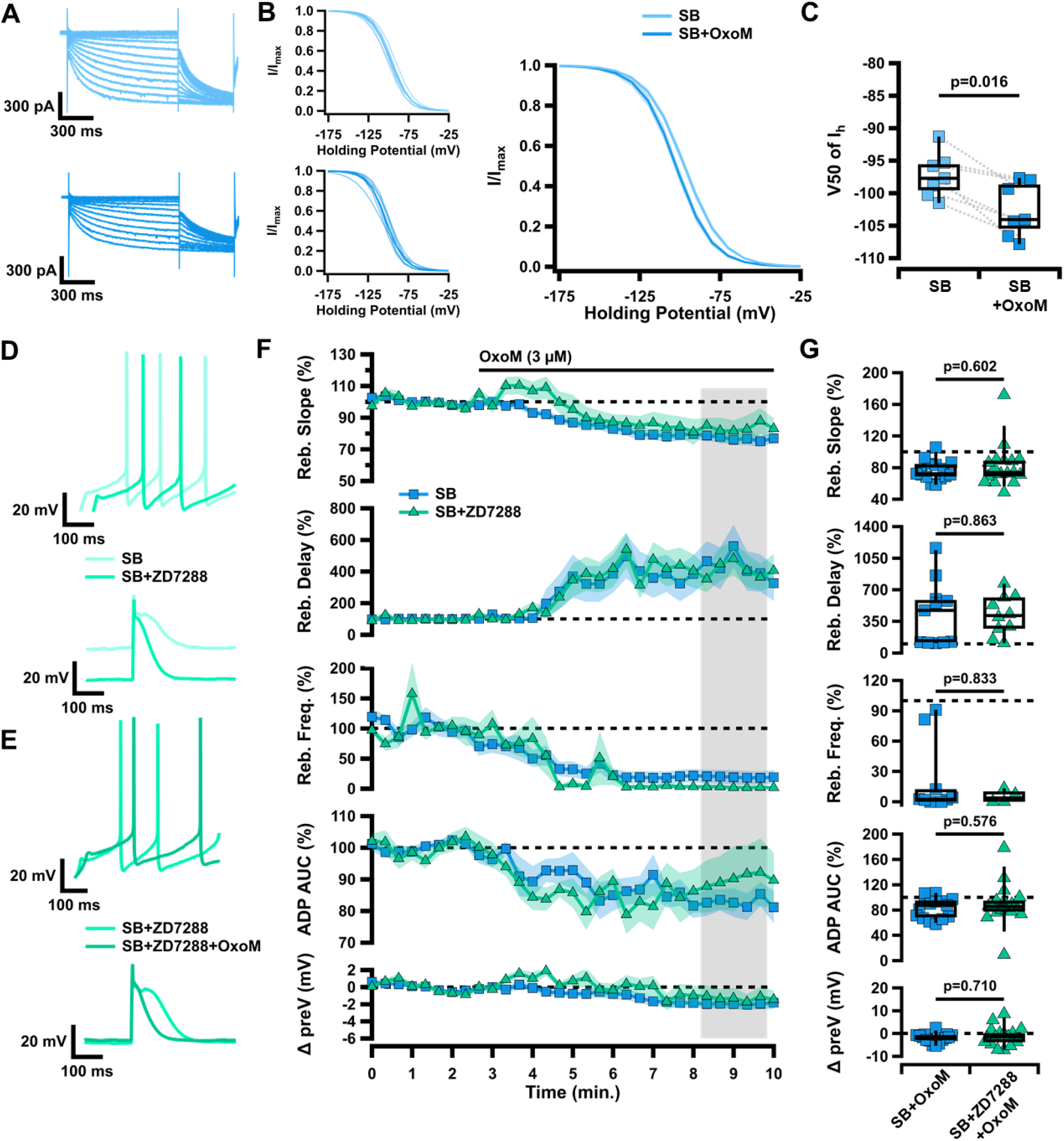
Changes in HCN channel activation are not responsible for muscarinic inhibition of rebound in ventral tier SNc neurons. **A**, Sample traces of HCN-mediated current measured in voltage clamp, in SB before (light blue) and after (dark blue) application of OxoM. **B**, Normalized activation curves of I_h_ tail current before (top left) and after (bottom left) application of OxoM, shown combined on right. Data presented as individual cells and their averages (left) and average ±SEM (right). **C**, Box plot showing V50 of I_h_ in SB before (light blue) and after (dark blue) application of OxoM. **D**, Sample traces of rebound (top) and the ADP (bottom) in SB before (light green) and after (green) application of ZD7288. **E**, Sample traces of rebound (top) and the ADP (bottom) in SB+ZD7288 before (green) and after (dark green) application of OxoM. **F**, Normalized rebound slope (top), rebound delay (second), rebound frequency (third), ADP AUC (fourth), and hyperpolarized baseline (bottom) as a function of time. Data presented as average ±SEM. **G**, Box plots representing individual cell averages in shaded regions of F.

Because the ventral tier, ADP-expressing SNc dopaminergic neurons also show larger I_h_ vs dorsal tier SNc neurons (Neuhoff et al., 2002), we hypothesized that OxoM selectively inhibits rebound in the ventral tier SNc because of its inhibition of I_h_. To test this, we applied the HCN channel blocker ZD7288 (ZD; 10 µM) prior to OxoM application. We found that OxoM-mediated inhibition of rebound was maintained even when HCN channels were blocked (Figure 4F-G). There was no significant difference between control and ZD conditions in the OxoM reduction of rebound slope (SB n=15, 76.34±3.30%; SB+ZD n=17, 82.17±6.64%; Wilcoxon rank-sum p=0.602), enhancement of rebound delay (SB n=11, 437.65±107.74%; SB+ZD n=10, 419.58±69.00%; Wilcoxon rank-sum p=0.863), or reduction in rebound frequency (SB n=10, 19.61±11.19%; SB+ZD n=4, 5.19±3.21%; Wilcoxon rank-sum p=0.833). These results demonstrate that although OxoM inhibits I_h_, this inhibition is not responsible for the muscarinic reduction of rebound activity. Further, there was no significant difference in the effect of OxoM on the ADP (SB n=15, 83.19±4.24%; SB+ZD n=17, 89.48±8.11%; Wilcoxon rank-sum p=0.576) or hyperpolarized baseline (SB n=15, -1.94±0.54 mV; SB+ZD n=17, -1.18±1.03 mV; Wilcoxon rank-sum p=0.710). Thus, here we establish that though OxoM shifts HCN activation to a lower membrane potential, this is not responsible for the OxoM effect on dopaminergic rebound activity.

### Simultaneous blockade of HCN and T-type calcium channels is not sufficient to occlude OxoM reduction of rebound

Previous research has shown synergistic activity between HCN channel activity and other intrinsic ion channels (Cobb-Lewis et al., 2023). Therefore, we hypothesized that HCN channels and TTCCs may together mediate the effects of OxoM on rebound. We performed experiments simultaneously blocking both channel types to determine if their cumulative effects occlude those of OxoM. We applied OxoM in the presence of TTA-P2 and ZD7288 and found that blocking both T-type current and I_h_ concurrently did not reduce the effect of OxoM on rebound activity (Figure 5 B-C). There was no difference between bath solution containing synaptic blockers alone or with TTA-P2 and ZD7288 on rebound slope (SB n=15, 437.65±107.74%; SB+TTA+ZD n=8, 66.76±6.54%; Wilcoxon rank-sum p=0.325) or hyperpolarized baseline (SB n=15, -1.94±0.54 mV; SB+TTA+ZD n=8, -1.18±1.03 mV; Wilcoxon rank-sum p=0.392). Because of the drastic effects of TTA-P2+ZD7288 alone on rebound (Figure 5D), we were unable to accurately measure the effect of OxoM on rebound delay and frequency, as the action potential timing slowed beyond what would typically be considered rebound firing when TTCCs and HCN channels are blocked. This created floor and ceiling effects that made the percent change measurement inadequate (Figure 5E). Because TTA-P2 also eliminated the ADP, we could not measure the effect of OxoM on ADP size. Therefore, we used rebound slope as the measure of rebound activity for this experiment. Together, these experiments show that neither TTCCs nor HCN channels, alone or in combination, mediate the effect of OxoM on dopaminergic rebound activity.

**Figure 5.**
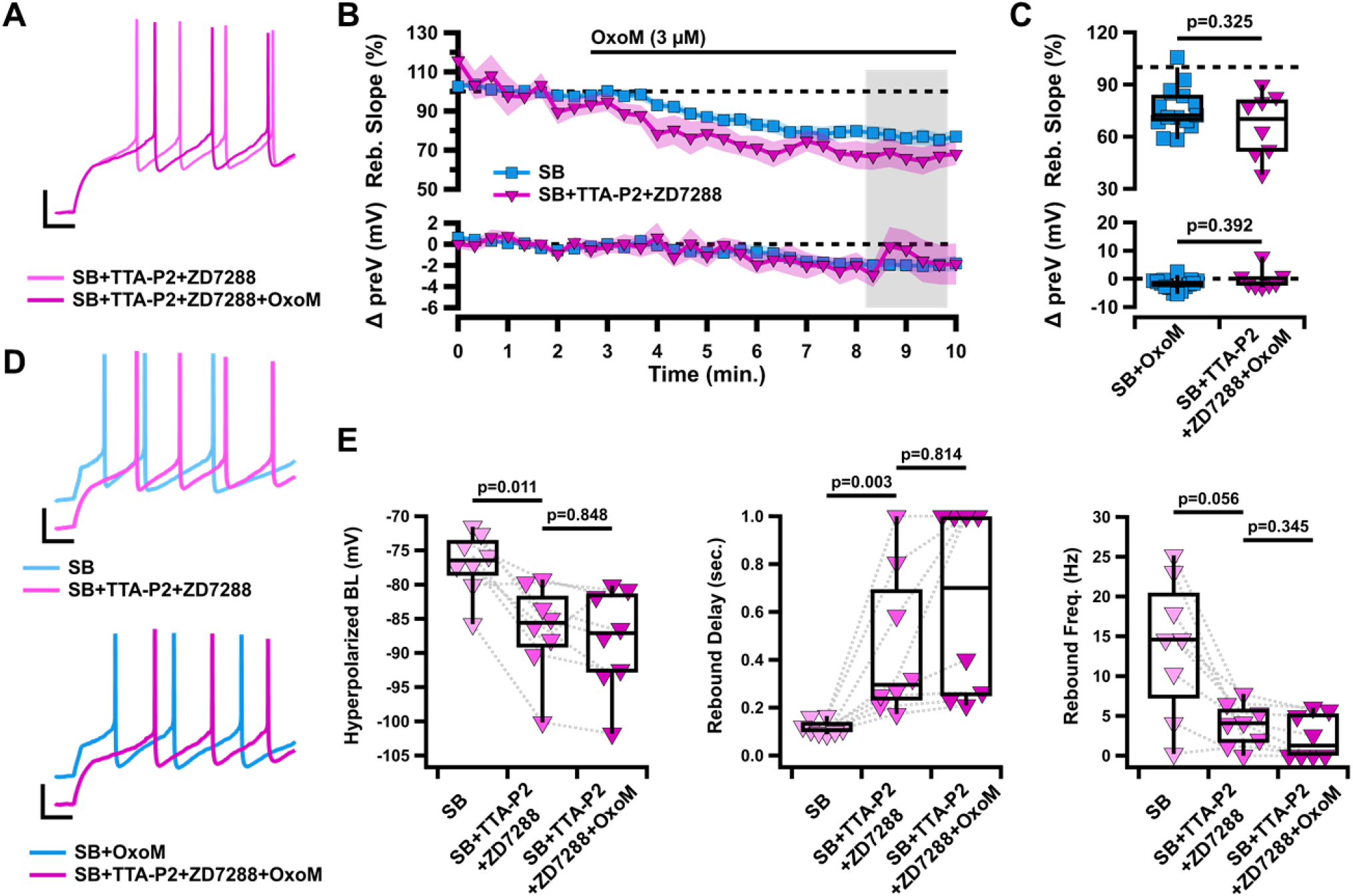
Simultaneous inhibition of T-type calcium and HCN channels does not occlude the effects of OxoM on rebound. **A**, Sample traces of rebound in SB+TTA-P2+ZD7288 (fuchsia), and SB+TTA-P2+ZD7288+OxoM (dark fuchsia). **B**, Normalized rebound slope (top) and hyperpolarized baseline (bottom) as a function of time. Data presented as average ±SEM. **C**, Box plots represent individual cell averages in shaded regions of B. **D**, Sample traces of rebound in SB (light blue) vs SB+TTA-P2+ZD7288 (fuchsia) (top) and SB+OxoM vs SB+TTA-P2+ZD7288+OxoM (dark fuchsia) (bottom). **E**, Box plots showing hyperpolarized baseline (left), rebound delay (middle), and rebound frequency (right) of the same cells in SB (light fuchsia), SB+TTA-P2+ZD7288 (fuchsia), and SB+TTA-P2+ZD7288+OxoM (dark fuchsia).

### Blocking A-type potassium channels enhances the effect of OxoM on rebound activity

Previous work has shown that the A-type potassium current (I_A_) also plays a role in modulating rebound firing in SNc neurons (Amendola et al., 2012; Tarfa et al., 2017). A-type potassium channels are activated when the cell depolarizes after a period of hyperpolarization. This slows the rebound depolarization and reduces rebound firing. I_A_ has an opposite influence from I_h_ and TTCCs on rebound activity, and blocking this current completely eliminates the rebound delay. In Figure 6, we applied OxoM in the presence of the A-type potassium channel blocker AmmTx3 (100 nM). Because AmmTx3 enhanced rebound firing so strongly, it was not possible to measure the rebound slope of the first rebound spike, as it occurred immediately upon release from hyperpolarization (Figure 6A, bottom). For this same reason we were unable to measure changes in rebound delay. Here, we instead measured the slope between the first (immediate) and second rebound action potentials and measured the rebound frequency of the first two spikes after release from hyperpolarization. We found that OxoM had an enhanced effect on rebound activity in the presence of AmmTx3 (Figure 6C-D). With A-type potassium channels blocked, OxoM more strongly reduced rebound slope than OxoM alone (SB n=15, 76.34±3.30%; SB+AmmTx3 n=7, 50.00±5.48%; Wilcoxon rank-sum p=0.002). However, in the presence of AmmTx3, OxoM decreased rebound frequency and ADP size to the same extent as when applied alone (rebound frequency: SB n=10, 19.61±11.19%; SB+AmmTx3 n=4, 4.86±0.51%; Wilcoxon rank-sum p=0.445; ADP size: SB n=15, 83.19±4.24%; SB+AmmTx3 n=7, 86.37±8.64%; Wilcoxon rank-sum p=0.945). The OxoM effect on the hyperpolarized baseline was also unchanged in the presence of AmmTx3 (SB n=15, - 1.94±0.54 mV; SB+AmmTx3 n=7, -2.55±0.63 mV; Wilcoxon rank-sum p=0.490). These experiments show that OxoM does not inhibit rebound by enhancing A-type potassium current activity, and supports the idea that muscarinic activation may actually inhibit I_A_ in dopaminergic neurons (Gantz and Bean, 2017).

**Figure 6.**
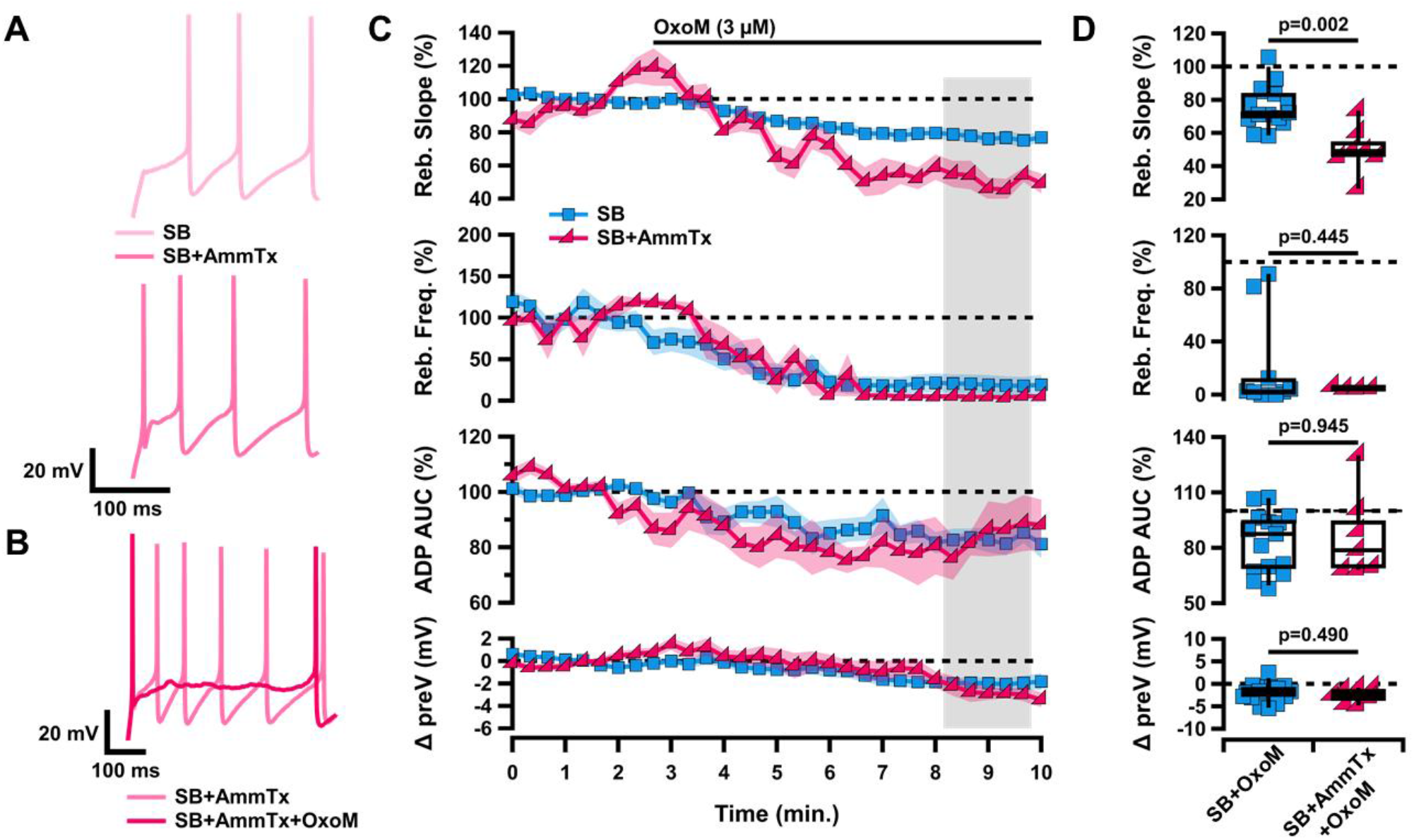
Muscarinic inhibition of rebound and the ADP of ventral tier SNc neurons is not mediated by A-type potassium channels. **A**, Sample traces of rebound in SB (light pink, top) and SB+AmmTx (pink, bottom). **B**, Sample traces of rebound in SB+AmmTx (pink) and SB+AmmTx+OxoM (dark pink). **C**, Normalized rebound slope (top), rebound frequency (second), ADP AUC (third), and hyperpolarized baseline (bottom) as a function of time. Data presented as average ±SEM. **D**, Box plots representing individual cell averages in shaded regions of C.

## Discussion

Here we found that muscarinic activation inhibits rebound in ventral tier SNc neurons more strongly than in dorsal tier SNc neurons. We found that this rebound inhibition is a direct result of mAChR activation and is not mediated through the modulation of pre-synaptic neurotransmitter release. Counter to our original hypotheses, we found that muscarinic activation does not inhibit rebound in SNc neurons by inhibiting T-type calcium channels or hyperpolarization activated cation channels either alone or in combination. Finally, we found that blocking A-type potassium channels enhanced muscarinic-mediated inhibition of rebound. Therefore, we conclude that mAChR activation inhibits rebound in SNc neurons through a non-canonical rebound mechanism.

The ventral and dorsal tier of the SNc differ in molecular markers (Poulin et al., 2014, 2020; Wu et al., 2019), electrophysiological characteristics (Neuhoff et al., 2002; Evans et al., 2017), and circuit connectivity (Evans et al., 2020). One prominent difference between these populations is their differential ability to rebound. The ventral tier neurons can be considered “rebound-ready” because of the strong expression of TTCCs and HCN channels (Neuhoff et al., 2002; Evans et al., 2017). Our finding that mAChR activation differentially affects the ADP-expressing (ventral) and non-ADP expressing (dorsal) SNc neurons is further evidence that these two populations process information in unique ways. Dopaminergic neuron rebound activity has been reported *in vivo* in primates and rodents (Fiorillo et al., 2013a, 2013b; Gut et al., 2022; Dong et al., 2024), and has been hypothesized to function as a safety or relief signal after an aversive stimulus (Wang and Tsien, 2011; Budygin et al., 2012; Fiorillo et al., 2013a; Lerner et al., 2015; Jong et al., 2019). The dynamic modulation of rebound activity by mAChRs is an important component in the acetylcholine-dopamine interactions that occur in the midbrain and can ultimately influence dopamine release in other brain structures, such as the striatum.

In physiological conditions, the main source of acetylcholine release in the SNc is from the cholinergic neurons of the pedunculopontine nucleus (PPN) (Clarke et al., 1987; Mena-Segovia et al., 2008; Dautan et al., 2016; Xiao et al., 2016; Estakhr et al., 2017), a brainstem structure that is involved in the coordination of movement and motor learning (Roseberry et al., 2016; Li and Spitzer, 2020; Dautan et al., 2021). Cholinergic axons have been identified in the SNc, particularly in the dendron bouquets specific to the ventral tier (Crittenden et al., 2016), and muscarinic receptor activation in the SNc is critical for PPN stimulation to generate long-lasting dopamine signals in the striatum (Forster and Blaha, 2003). Future work is needed to fully dissect the influence of endogenous acetylcholine released from the PPN onto the SNc, and to determine whether M5 muscarinic receptor activation reduces intrinsic dopamine rebound activity *in vivo*.

Rebound activity in dopaminergic neurons is controlled by three main channels: T-type calcium channels, hyperpolarization-activated cation channels, and A-type potassium channels (Neuhoff et al., 2002; Amendola et al., 2012; Evans et al., 2017; Tarfa et al., 2017). The SNc receives strong inhibitory input from multiple basal ganglia nuclei (Saitoh et al., 2004; McGregor et al., 2019; Evans, 2022; Gut et al., 2022) which hyperpolarize the membrane and recruit these cation channels. Previous work has shown that G_q_-coupled muscarinic receptors can inhibit TTCCs and do so particularly strongly for the CaV3.3 TTCC subtype in cultured cells (Hildebrand et al., 2007). Previous work has also shown that muscarinic receptor activation inhibits HCN channels in striatal cholinergic neurons (Zhao et al., 2016), but enhances it in vestibular ganglion neurons (Bronson and Kalluri, 2023). Here we show that in SNc dopaminergic neurons, mAChR activation inhibits HCN activity (Figure 4). Together, these results suggest that M5 receptor activation would reduce rebound activity through TTCCs and HCN channels. Surprisingly, we found that the inhibition of these channels was not necessary for muscarinic receptor activation to inhibit rebound in SNc dopaminergic neurons. On the other hand, previous work has shown that G_q_-coupled receptors inhibit A-type potassium channels in dissociated dopaminergic neurons (Gantz and Bean, 2017). Because A-type activation reduces rebound activity, this result suggests that M5 activation would enhance rebound by inhibiting A-type channels. However, we see that M5 activation reduces rebound. Therefore, our findings support the idea that M5 activation causes multiple distinct physiological changes in SNc neurons to both positively and negatively influence rebound.

Previous work has found that brief (seconds to minutes) application of OxoM to brain slices increases neural firing and somatic calcium in dopaminergic neurons (Gronier and Rasmussen, 1998; Foster et al., 2014). By contrast, our experiments did not show a significant OxoM effect on tonic firing of either ADP-expressing or non-ADP-expressing SNc neurons. This discrepancy may be due to the difference in timing of the OxoM application (short vs long exposure), the difference in OxoM concentration (10 µM vs 3 µM), or the different electrophysiological technique used (perforated-patch vs whole-cell). Interestingly, another study found that transient and long-lasting muscarinic stimulation caused opposing neural responses in dopaminergic neurons (Fiorillo and Williams, 2000). These studies highlight the complex interactions between acetylcholine and dopamine in the midbrain.

Together our findings reveal a previously unknown acetylcholine-dopamine interaction that occurs in the midbrain. The selective inhibition of rebound in the vulnerable SNc subpopulation by muscarinic receptor activation is important for our understanding of the complex interplay between the dopaminergic and cholinergic systems of the healthy brain. Because the dopaminergic neurons of the SNc and their cholinergic inputs from the brainstem degenerate in Parkinson’s disease (Yamada et al., 1990; Rinne et al., 2008; Sébille et al., 2019), the endogenous activation of M5 muscarinic receptors on SNc neurons is likely to be disrupted in this disorder. Future experiments will be critical for understanding how acetylcholine-dopamine interactions in the midbrain are altered in pathological conditions.

## Acknowledgements

This work was supported by the American Parkinson’s Disease Association Research Grant 2021APDA00RG00000209666, Parkinson’s Foundation Stanley Fahn Junior Faculty Award #PF-SF-JFA-1040267, and BRAIN Initiative K99/R00 award #R00NS112417 awarded to RCE; and by a National Institute of General Medical Sciences T32 predoctoral fellowship GM142520 awarded to MLB. We thank Dr. John Partridge and members of the Evans Lab for feedback on earlier versions of this manuscript.

